# Naturalistic face-directed attention predicts face recognition ability

**DOI:** 10.64898/2026.09.30.755511

**Authors:** David White, Victor P. L. Varela, Monique Piggott, James D. Dunn, Sebastien Miellet, Alice Towler, Bojana Popovic

**Author notes:** Corresponding Author: David White, School of Psychology, Mathews Building, UNSW Sydney, Kensington, NSW 2052, Australia.

## Abstract

Recognising people precedes and informs most social interactions, yet it is not clear why individuals differ so markedly in face recognition ability. Exposure-based accounts propose that it is shaped by visual experience with faces—known as ‘face diet’—but most prior work treats exposure as passively acquired, thereby masking the active and dynamic nature of visual attention in natural settings. Here we examined whether spontaneous attention to faces in everyday environments predicts face recognition ability. Eighty participants wore mobile eye-tracking glasses while walking through a busy university campus and during a face-to-face interaction, before completing standard measures of face recognition ability. Fixations directed to faces showed stable individual differences in both settings. People who looked more at faces scored higher on our composite measure of face recognition ability, and gaze in the two settings predicted ability independently, together accounting for 22% of its variance. These findings suggest that differences in how people actively sample visual information from their environments may be an overlooked source of variance in perceptual expertise.

## INTRODUCTION

Recognising faces underpins everyday social behaviour. Yet face recognition ability varies strikingly across the population (White & Burton, 2022)—from people who have difficulty recognising close friends and family (developmental prosopagnosia: Duchaine & Nakayama, 2006; Barton & Corrow, 2016) to those with extremely high face recognition ability (super-recognisers: Ramon et al., 2019; White, 2026). These differences can have a profound impact on social lives, with lower ability associated with interpersonal difficulties (Dalrymple et al., 2014; Yardley et al., 2008) and are strongly implicated in general disorders of social cognition (Minio-Paluello et al., 2020). They also have societal implications, for example in selecting people suited to verifying photo-ID (White et al. 2014) or in forensic settings (Dunn et al., 2023).

The biological basis of perceptual expertise with faces is not understood. Twin studies show that face recognition ability is strongly heritable, with shared environment accounting for little additional variance (Wilmer et al., 2010; Shakeshaft & Plomin, 2015), leading some researchers to argue that experience plays a limited role in its development (McKone et al., 2012). However, face recognition ability continues to improve into mid-adulthood (Germine, et al., 2011; Dunn et al., 2020), indicating substantial experience-dependent change. Resolving this apparent tension requires a theory of how visual experience with faces—often conceptualised as a “face diet”—accounts for individual differences in face recognition ability (Pickron & Bayet, 2026).

At the group level, differences in face diet are thought to underlie effects such as the other-race effect, whereby people are typically better at recognising faces from their own race than others (Meissner & Brigham, 2001). However, this focus on group-level exposure could mask substantial individual variation in the visual environments people experience. Wearable camera recordings reveal substantial variability in the quantity and quality of faces present in everyday environments (Oruc et al., 2019; Sugden et al., 2014), and self-reported differences in exposure to people from other ethnic groups can explain some variation in expertise with other-race faces (Chiroro & Valentine, 1995; McKone et al., 2019).

Whether variation in everyday face diet can account for individual differences in face recognition ability remains unclear. This may be because face diet is treated as passive exposure, rather than actively constructed by the viewer. For example, childhood hometown size is only weakly and inconsistently related to face recognition ability across studies (Balas & Saville, 2015, 2017; Sunday, et al., 2019; Andrews et al., 2026). Recent innovative work has begun to sample visual exposure to faces in the environment using wearable cameras—with some preliminary evidence of association between the amount of faces in the environment and face recognition ability in a small group of adults with Autism Spectrum Disorder (Kamensek, Iarocci & Oruc, 2024). However, these studies do not measure how people actively sample face information encountered in their environment.

On the other hand, social attention research has shown that when participants view visual arrays and scenes involving people on computer screens, there are large individual differences in the extent to which people attend to faces. These differences emerge early in life and twin studies show strong genetic influence from 5 months (Constantino et al. 2017; Portugal et al., 2024, 2025; Richmond et al., 2017). This raises the possibility that genetic differences in social attention may shape the environments individuals experience, which in turn may contribute to differences in perceptual abilities (Scarr & McCartney, 1983). Screen-based studies also show that adults with developmental prosopagnosia spend less time fixating on faces in images of social scenes (Bobak et al., 2016), and super-recognisers spend more time (Linka et al., 2022; but see de Haas et al., 2019).

Here we test the association between face recognition ability and active visual sampling by measuring individuals’ spontaneous attention to faces in natural social settings—moving beyond screen-based eye tracking to dynamic and interactive social environments. Attention to people that are physically present differs substantially from viewing images and varies with social context (Risko et al., 2016; Peterson et al., 2016) and so testing this in naturalistic settings provides a critical test of environmental influences on face recognition ability. To measure face-directed attention ‘in the wild’ we use a novel approach developed in our prior work—combining wearable eye tracking with person detection software to automatically map fixations made towards people in the environment (Varela et al., 2023; see also Hessels et al., 2020). In addition to establishing a link between face perception ability and social attention in natural settings, our results show the potential for this reproducible method for studying social attention more broadly.

## METHODS

### Transparency and openness

De-identified participant-level behavioural, self-report, and eye-tracking summary measures are publicly available (ANONYMOUS REVIEW LINK: https://osf.io/mu9gc/overview?view_only=a74a73fd70874cebb69a9a6286645e20). Raw scene videos can not be shared because they contain identifiable bystanders recorded on a public university campus. Code for the processing pipeline is publicly available (ANONYMOUS REVIEW LINK: https://anonymous.4open.science/r/natural-face-attention-F4B9/README.md), including sample Pupil Cloud and OpenPose outputs to enable reproduction of the social-attention measures.

### Participants

Participants were 79 psychology students from the University of New South Wales (UNSW), and 9 members of the public recruited from the UNSW Face Research Registry. Registry participants were recruited to provide a broader sampling of face recognition ability, with each achieving an average score on three standard face tests that placed them at least 1.7 standard deviations above the mean score (CFMT+, Russell et al. 2009; Glasgow Face Matching Test, Burton et al., 2010; UNSW Face Test, Dunn et al., 2020). This is a lower criteria compared to prior studies selecting ‘super-recognisers’ (e.g. Dunn et al., 2025). We excluded eight student participants due to corrupt eye-tracking data (n = 5) or because they did not complete the experiment (n = 3). Student participants received course credit and participants from our research registry received a small payment. After exclusions, analysis was performed on a sample of 80 participants (42 female, 33 male, 5 non-binary) ranging from ages 18 to 63 years old (M = 23.2 years, SD = 9.81). All participants had normal or corrected to normal vision.

### Procedure

Participants completed a standardised experimental procedure in naturalistic environment while their eye movements were measured using wearable eye-trackers. The first phase was the ‘campus walk’. This consisted of two loops around the same pre-defined route in a busy university campus during term time, and we conducted the tests between 9am and 5pm so that the number of people on campus was comparable for all participants. This route was explained both verbally and visually to the participant prior to the walk commencing, and before calibrating the wearable eye-tracker according to manufactiurer guidelines. During the walk, the experimenter followed the participant to prevent any issues with directions as well as to measure lighting levels. At the end of the walk, the participant returned to the testing room. The recording time for the campus walk was 13 minutes on average.

The ‘face-to-face interaction’ phase of the experiment began once the participant had returned to the testing room. They were asked to stand 1.5m in front of the experimenter who faced them directly in a corridor. The experimenter then engaged in a structured conversation with the participant. This conversation involved the experimenter explaining details of the study and encouraging any questions or comments from the participant. Overall, the conversation allocated equal time for the participant to both speak and listen, with the average face-to-face interaction recording lasting 1 minute.

After this interaction was complete, participants were asked to remove the eye tracking glasses. For the final part of the study, all participants completed the face recognition ability measures which took approximately 20 minutes in total. University student participants were allocated a longer study session and also completed additional individual differences measures which took an additional 20 minutes on average. Details of these measures and the eye-tracking equipment and data processing are provided below.

### Face recognition ability measures

We measured face recognition ability using an objective laboratory measure of face memory accuracy (Cambridge Face Memory Test Long Form, CFMT+; Russell et al., 2009) and a self-report measure intended to capture participants’ everyday face recognition experience outside the lab (Prosopagnosia Index, PI-20; Shah et al., 2015). To obtain an overall index of face recognition ability combining lab-based measurement and participanty’s everyday experience (see Burns et al., 2023) we computed a composite score as the mean of CFMT+ and PI-20 z-scores from normative scores reported in prior work (Bobak, Pampoulov & Bate, 2016; Shah et al., 2015).

The CFMT+ is an extended, more difficult version of the CFMT (Duchaine & Nakayama, 2006) designed to improve discrimination among higher-ability individuals. Participants study sets of faces and are subsequently tested by selecting learned identities from arrays containing foil faces; performance is scored as total correct out of 102, with higher scores indicating better face memory. The PI-20 is a 20-item self-report questionnaire assessing perceived face recognition difficulties. Participants rate each item on a 5-point Likert scale (1 = strongly disagree, 5 = strongly agree), and responses are summed to yield a total score out of 100, with higher scores indicating poorer self-perceived face recognition ability. PI-20 z-scores were reverse-scored prior to averaging so that higher values on the composite reflected better face recognition ability across both measures.

### Individual difference measures

We selected individual difference measures for inclusion in the study where prior work had shown either an association with facial recognition ability and social attention behaviour. We measured social anxiety using Liebowitz Social Phobia Scale (LSAS; Liebowitz, 1987), extraversion using the extraversion-introversion subscale of the Eysenck Personality Inventory (EPI; Eysenck & Eysenck, 1963) and autism-like traits using the 28-item Autism Quotient Short Form (AQSF; Hoekstra et al., 2011). We also assessed state anxiety using subscale form Y-1 from State-Trait Anxiety Inventory (STAI-20; Spielberger et al., 1983). Although intercorrelations between these measures were consistent with literature on anxiety and trait autism, we did not find reliable associations between these measures and either face recognition ability or social attention behaviour. Methodlogical details and full reporting of associations between these and all of our other measures are in Supplementary Materials (Table S1).

### Eye-Tracking and data processing

Participants eye gaze was recorded using the Pupil Labs Invisible wearable eye tracking glasses (Tonsen et al., 2020). These devices recorded videos of participants field of view using a standard camera (30Hz, 1088 x 1080 pixels, covering full range of view) and eye gaze coordinates via infa-red eye cameras (200Hz @ 192 x 192 pixels) to a smartphone via USB cable. Raw gaze samples were classified into fixations and saccades using the default settings in manufacturer’s cloud processing (Pupil Cloud v5.6 Fixation Detection; maximum dispersion = 1.5° visual angle, minimum duration = 60 ms) and exported as pixel coordinates in the scene (i.e. frontal camera frame).

To quantify attention to people, fixation coordinates were mapped onto person landmarks detected in each video frame using OpenPose (Cao et al., 2021), following the same approach as Varela et al. (2023). OpenPose provided pixel coordinates for 5 head landmarks and 20 body landmarks (7 torso, 12 legs) for each detected person. Fixations to faces were defined as any fixation within 70 pixels of a head landmark, and fixations to bodies as any fixation within 70 pixels of a body landmark. When a fixation fell within 70 pixels of both head and body landmarks, it was assigned to the region containing the closest landmark. Proportions of fixations to faces and bodies were computed as a proportion of total fixations, calculated only for frames in which a fixation was detected. Code used to compute social attention measures from wearable eye-tracking data files is avaliable via the GitHub link above.

## RESULTS

Figure 1 shows the proportion of fixations to faces and bodies separately for when participants walked through campus and during a face-to-face interaction. Proportion of fixations made to passers-by in the campus walk were consistent with a prior study that developed the methods reported here (Face fixations: M = 0.061; SD = 0.035; Body fixations: M = 0.123; SD = 0.070; see also Varela et al., 2023), but was substantially lower than during face-to-face interaction with the experimenter (Face fixations: M = 0.665; SD = 0.147; Body fixations: M = 0.054; SD = 0.061).

**Figure 1.**
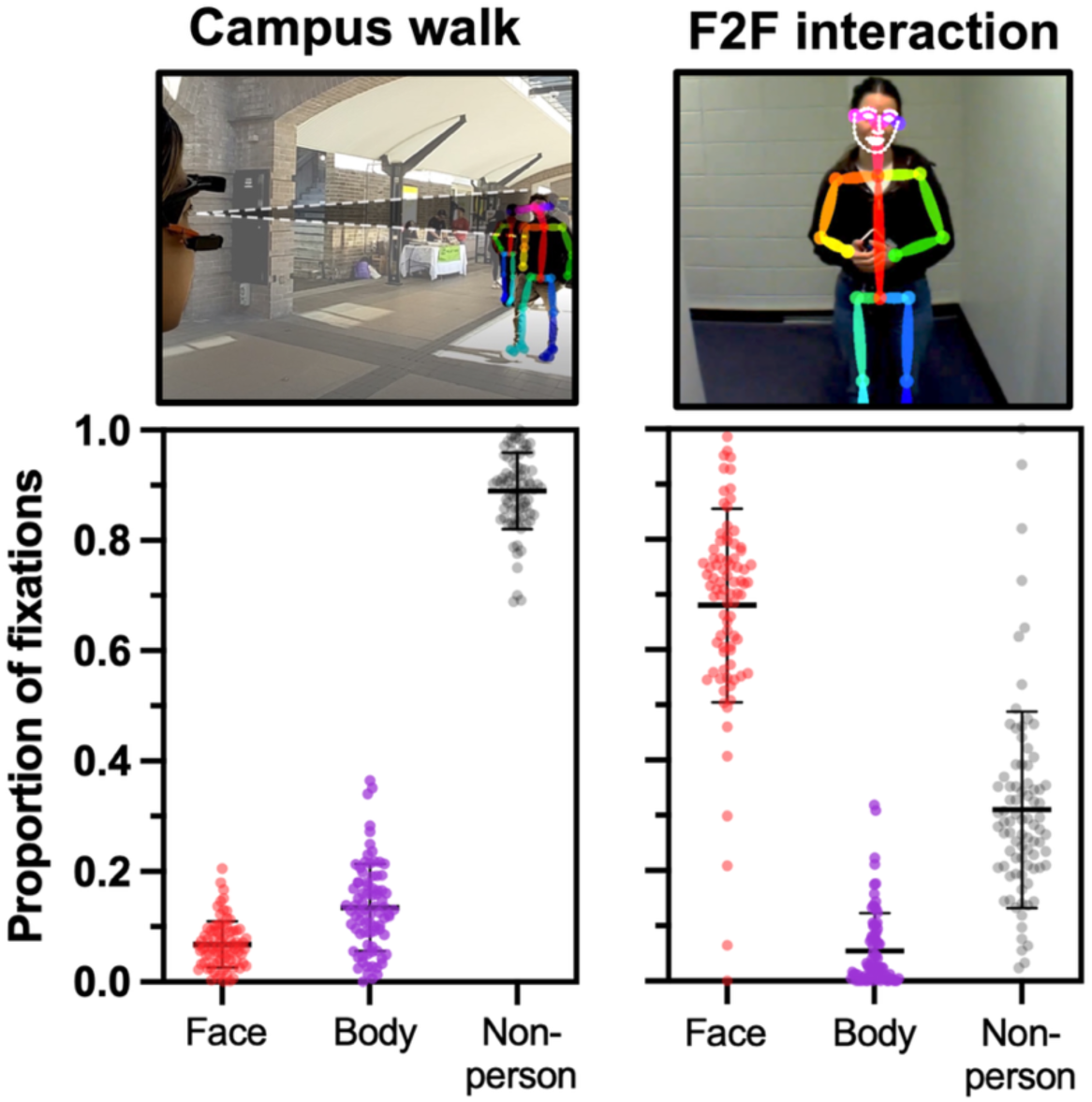
Gaze directed to faces and bodies in two phases of the experiment. Top panels show OpenPose person detection applied to wearable eye tracking devices in two naturalistic scenarios. An example video of naturalistic social attention measurement is available via OSF and code is available via GitHub (see Methods: Transparency and openness). Bottom panels show distributions of participants face, body and non-person directed fixations for the non-interactive campus walk (left) and interactive F2F scenario (right). Data points show the proportion of fixations for individual participants, with horizontal black lines showing the median and 95% CIs. The people shown in this figure are study authors.

Figure 1 also shows wide individual differences in proportion of fixations to faces and bodies in both phases of the study. This is especially noticeable for the face-to-face interaction, where some individuals barely fixated on the experimenter’s face at all—while others maintained fixation on their face throughout. Split-half reliability based on Pearson’s r showed trait-like stability in these individual differences (Walk face gaze *r* = 0.815; Walk body gaze *r* = 0.827; F2F face gaze *r* = 0.584; F2F body gaze *r* = 0.627).

Figure 2 shows the associations between face recognition ability and gaze toward faces and bodies. We found reliable correlations between proportion of gaze to faces and face recognition ability (Figure 2, top panels), in both the campus walk, rho (78) = .357, 95% bootstrap CI [.147, .561], p = .001, and the F2F interaction phase, rho (78) = .349, CI [.137, .543], p = .002. Gaze to bodies (Figure 2, bottom panels) was unrelated to ability during the F2F interaction, rho (78) = .007, CI [−.222, .224], p = .950, and only marginally related during the campus walk, rho (78) = .230, CI [.001, .439], p = .040.

**Figure 2.**
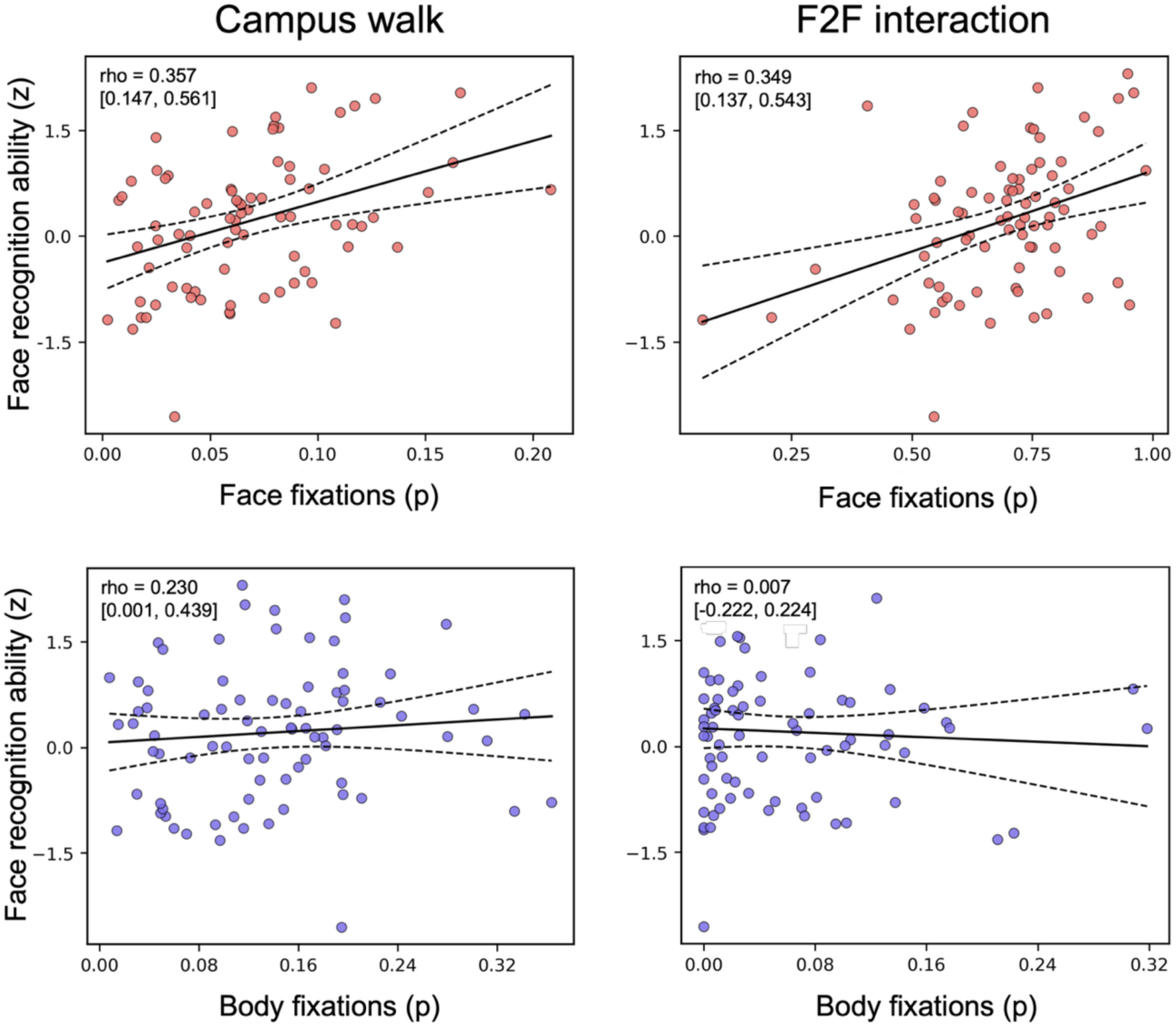
Association between face recognition ability and social attention in two naturalistic settings. Scatterplots on the top show proportion of fixations to faces as a function of face recognition ability in campus walk (left) and face-to-face interaction (right). Scatterplots on the bottom show proportion of fixations to bodies in these same contexts. Best fitting regression lines are shown with 95% confidence bands denoted by dashed lines.

Face-directed gaze in the two contexts was only weakly correlated, rho (78) = .219, CI [−.008, .430], p = .051, so we also entered both as simultaneous predictors of face recognition ability. The model explained 22.4% of the variance, R² = .224, 95% CI [.101, .391], adjusted R² = .204, F(2, 77) = 11.13, p < .001, with both contexts contributing independently (campus walk β = .286, CI [.083, .490], p = .007; F2F interaction β = .311, CI [.107, .515], p = .004).

## DISCUSSION

Across two everyday scenarios, we found wide individual variation in viewer’s attention to faces and people. These differences are consistent with lab-based studies showing substantial heritability in social attention (e.g., Constantino et al., 2017; Portugal, et al., 2024), confirming that these differences are observed with relative stability even in more unconstrained natural settings. This is the first study of this scale that has used reproducible measures of these individual differences in natural settings, and so it will of be important to extend these findings to different social settings and contexts in future. Nevertheless, our study underlines the clear potential for using automated detection software combined with wearable eye-tracking to study visual attention in these settings (Culham & Deligiannis, 2026).

Our results also have implications for the study of perceptual expertise. Face perception is often used as a window to examine how cognitive traits emerge from environmental constraints. These constraints are typically framed in terms of the face information available in the environment and the operations performed on it (Young, 2017), and yet exposure in the environment is very rarely measured. Some recent work has begun to measure naturalistic “face diet” using wearable cameras (Oruc et al. 2019; Sugden et al, 2014), but this only provides exposure statistics to faces present in the environment—and not the faces that are attend to. In our study, even within the brief window of experience we captured, individuals differed markedly in how much face information they actively acquired from broadly the same environments. This suggests that the face diet most relevant to face expertise is not passively received—but actively constructed through our idiosyncratic tendency to fixate faces.

We found that the amount of attention directed to faces predicted individuals’ face recognition ability. This relationship was not consistently observed for body-directed attention, indicating that the effect is specific to faces rather than a general bias towards people—raising the possibility that face recognition ability may, in part, reflect downstream consequences of individual differences in social attention. Differences in attention to faces show substantial heritability even at an early age (e.g., Constantino et al., 2017), whereas face recognition ability follows a more protracted developmental trajectory (Germine et al., 2011) and may show weaker or less stable heritability earlier in life (Shakeshaft & Plomin, 2015; Wilmer et al., 2010; cf. Zhu et al., 2010). Because variation in social attention appears to determine visual input available for learning about faces, it is likely that the heritability of face recognition ability is related to early variation in social attention, but this would need to be confirmed in developmental studies.

Notably, the association between face-directed attention and face recognition ability emerged despite limited opportunities for participants to shape their environment within the experiment: faces were unfamiliar prior to the study, the route was pre-determined, and social interaction was semi-scripted. In everyday life, individuals vary widely in their social environments, including the number of people they know (Dunbar, 2020; Jenkins, 2018), the nature of their relationships (Engfors et al., 2024), the proportion of familiar faces they encounter (Kamensek et al., 2024; Oruc et al., 2019) and the distance they choose to view faces from (Kamensek et al., 2024). This multidimensionality entails that exposure is fundamentally idiosyncratic to individual viewers (Burton, 2026). We have shown that individual differences in social attention contribute meaningfully to exposure at the level of individuals, and face recognition ability. A deeper understanding of environmental influences on face recognition ability would require extending our investigation to less constrained settings, for example incorporating interactions with familiar individuals in daily life (Kamensek et al. 2024; Young et al., 1985; Young & Burton, 2018).

In summary our findings show that, outside the lab, individuals differ substantially in how they actively sample visual information, and that these differences are related to perceptual ability on lab-based tests. But our study samples only a small part of what is now discoverable about real-world perception (Culham & Deligiannis, 2026). Naturalistic measurement of perceptual input could have benefits across other fields of psychology, and for the development of computational models that are trained on naturalistic input (Phillips & White, 2026). Advances in wearable eye tracking and automated scene analysis now make it possible to measure active visual sampling at scale (Vong et al., 2024), providing new opportunities to understand how social, cognitive and perceptual abilities emerge from the actively constructed visual environments of individual viewers.

