## Supplementary Materials for "Naturalistic face-directed attention predicts face recognition ability"

### Individual differences measures

***State-Trait Anxiety Inventory (STAI) subscale form Y-1 (Spielberger et al., 1983).*** Participants completed the 20-item STAI-S (state anxiety), responding to each item using a 4-point scale ranging from "Not at All" to "Very Much So". Responses for all items are summed to calculate an overall score (/80) with higher scores indicating greater state anxiety.

***Liebowitz Social Phobia Scale (LSAS).*** The LSAS (Liebowitz, 1987) is a 24-item self-report questionnaire which assesses individuals' fear and avoidance towards social and performance-based scenarios. Two questions are posed for each scenario: the first question measures how anxious or fearful respondents feel towards the described situation, rated on a 4-point Likert scale (0 = none, 3 = severe) whilst the second question assesses how often respondents avoid a described situation, also rated on a 4-point Likert scale (0 = never, 3 = usually). Total scores are added together to calculate a final score out of 144. A higher score suggests more severe social phobia.

***Eysenck Personality Inventory (EPI).*** This study used only the extraversion-introversion subscale of the EPI (Eysenck & Eysenck, 1964) which contains 24 items aimed at measuring levels of extroversion. Participants respond with either "Yes" or "No" to each item and a total score is calculated by summing all 24 responses such that a score closer to 24 indicates greater levels of extraversion. This questionnaire also includes an additional 9-items which reflect a lie score, that is, a measure of how socially desirable participants are trying to be with their responses. A lie score equal to or greater than 5 indicates social desirability in participant responses.

***Autism Quotient Short Form (AQ-SF).*** The AQ-S (Hoekstra et al., 2011) is a 28-item self-report questionnaire (adapted from the 50-item Autism Quotient; Baron-Cohen et al., 2001) measuring the presence of autism spectrum symptoms within individuals. Participants respond on a Likert-scale (0= definitely agree, slightly agree, 1= slightly disagree, definitely disagree). Total scores are added together to calculate a final score out of 25. A higher score indicates higher levels of autism spectrum symptoms.

***Supplementary analysis of individual differences measures***. Full table of correlations between face recognition ability and each of the individual difference measures are shown in Table S1. The observed pattern of correlations between individual differences measures was consistent with existing literature. Extraversion was negatively associated with both social anxiety ($r = -.378, p < .01$) and trait autism ($r = -.665, p < .001$), aligning with extensive evidence linking lower extraversion (i.e., reduced sociability and positive affect) to higher levels of social anxiety and autism-related characteristics (Kaplan et al., 2015; Lodi-Smith et al., 2019; Lyon et al., 2021; Shirayama et al., 2022). Additionally, trait autism was positively associated with both social anxiety ($r = .452, p < .001$) and state anxiety ($r = .324, p < .01$), aligning with literature documenting the co-occurrence of autism with social anxiety disorder (see Montaser et al., 2023 for review), as well as heightened anxiety (Capp et al., 2025; Hwang et al., 2020). As reported in the main paper, despite these stable patterns of trait associations in individual difference measures, we did not observe reliable associations between these measures and face recognition ability or social attention.

| | Fix face (walk) | Fix face (F2F) | Fix body (walk) | Fix body (F2F) | CFMT | PI20 | LSAS | EPI-Extraversion | AQSF | STAI-20 |
|---|---|---|---|---|---|---|---|---|---|---|
| **Fixations face (walk)** | — | 78 | 78 | 78 | 78 | 77 | 69 | 69 | 69 | 69 |
| **Fixations face (F2F)** | 0.219 | — | 78 | 78 | 78 | 77 | 69 | 69 | 69 | 69 |
| **Fixations body (walk)** | 0.765*** | 0.142 | — | 78 | 78 | 77 | 69 | 69 | 69 | 69 |
| **Fixations body (F2F)** | 0.097 | -0.205 | 0.282* | — | 78 | 77 | 69 | 69 | 69 | 69 |
| **CFMT** | 0.288** | 0.298** | 0.263* | 0.127 | — | 77 | 69 | 69 | 69 | 69 |
| **PI20** | -0.322** | -0.326** | -0.146 | 0.149 | -0.436*** | — | 69 | 69 | 69 | 69 |
| **LSAS** | -0.146 | -0.128 | -0.009 | -0.059 | -0.025 | 0.235* | — | 69 | 69 | 69 |
| **EPI-Extraversion** | 0.231 | 0.118 | 0.211 | 0.092 | -0.088 | -0.027 | -0.378** | — | 69 | 69 |
| **AQSF** | -0.150 | -0.076 | -0.182 | -0.055 | -0.009 | 0.160 | 0.452*** | -0.665*** | — | 69 |
| **STAI-20** | -0.203 | -0.240* | -0.033 | -0.085 | -0.127 | 0.262* | 0.524*** | -0.165 | 0.324** | — |

**Table S1. Pairwise correlations among all measures.** The lower-left triangle reports Spearman's rank-order correlations (ρ), dashes indicate the diagonal, and the upper-right triangle reports the corresponding degrees of freedom (df). Significance is denoted by asterisks (* $p < .05$, ** $p < .01$, *** $p < .001$; two-tailed). Degrees of freedom vary across pairs due to missing data as only a subset of participants completed the individual differences tests (LSAS, EPI, AQSF, STAI-20)
